# Functional interplay between (p)ppGpp and RNAP in *Acinetobacter baumannii*

**DOI:** 10.1101/2025.07.04.663200

**Authors:** Anthony Perrier, Aurélie Budin-Verneuil, Régis Hallez

**Affiliations:** Bacterial Cell cycle & Development (BCcD), Biology of Microorganisms Research Unit (URBM), Namur Research Institute for Life Science (NARILIS), University of Namur, Namur (5000), Belgium; Université de Caen Normandie, CBSA UR 4312, Caen, France

**Keywords:** (p)ppGpp, stress response, RNAP, hydrophilin, desiccation, surface motility

## Abstract

The (p)ppGpp-dependent stress response is required for pathogenic bacteria to survive both outside and inside the host but the mechanisms behind this survival are mostly unknown. In this study, we characterize the (p)ppGpp metabolism in the opportunistic pathogen multi-drug-resistant *Acinetobacter baumannii*. We show that two stressful conditions potentially encountered during infection – iron starvation and polymyxin exposure – induce (p)ppGpp production. The absence of (p)ppGpp led to multiple consequences on the physiology of *A. baumannii*, including an increase of surface motility, a decrease in catalase activity, a poor survival upon nutrient starvation, a rapid killing during desiccation and a strong attenuation in a *Galleria mellonella* model of infection. Using a motility suppressor screen, we isolated multiple independent stringent mutations in *rpoB* and *rpoC* that suppress the (p)ppGpp-dependent phenotypes. By combining the suppressor screen with deep sequencing, we isolated dozens of additional mutants, expanding the list of putative stringent RNAP mutations described so far. Furthermore, our transcriptomic data reveal that (p)ppGpp deeply impacts the transcriptional landscape of *A. baumannii* on solid surface to induce many stress-related genes, including catalase and hydrophilins critical for tolerance to desiccation. This work highlights the functional interplay between (p)ppGpp and RNAP in the successful survival of *A. baumannii* in the environment but also during infection.

## Introduction

The stringent response (SR) is one of the mechanisms that facilitates bacterial adaptation to harmful conditions. In 1969, Cashel and Gallant identified two compounds related to the SR, the guanosine tetra-phosphate (ppGpp) and penta-phosphate (pppGpp) collectively referred to as (p)ppGpp (1). The SR was first referred to the growth arrest of *Escherichia coli* cells experiencing amino acid starvation due to increasing intracellular levels of (p)ppGpp. Now, the SR refers to any response caused by elevated levels of (p)ppGpp by any means (2). The intracellular levels of (p)ppGpp are mainly regulated by enzymes of the RSH family (RelA/SpoT Homologue). Most of bacteria possess one long bifunctional RSH, whereas the β- and γ-Proteobacteria harbor two long RSH proteins, one bifunctional (SpoT) and one monofunctional (RelA). Interestingly, based on *in silico* analyses and experimental evidence, the Moraxellaceae family to which belongs the genus *Acinetobacter* seems to have two long monofunctional RSH, RelA and SpoT displaying respectively functional synthetase and hydrolase activities (3,4). In addition to long RSH, some bacteria also possess short monodomain RSH with either a synthase or hydrolase activity, respectively called Small Alarmone Synthetase (SAS) or Hydrolase (SAH).

Since its discovery more than 50 years ago it has been often hypothesized that (p)ppGpp directly interacts with the bacterial RNA polymerase (RNAP). But it is only during the last decade that substantial progress has been made and two (p)ppGpp binding site (respectively referred to as site 1 and site 2) were mapped on the *E. coli* RNAP. The site 1 is located at the interface between the ω and β’ subunits (5–7) while the site 2 is created by the interaction of the transcription factor DksA with the β’ subunit (8). The position of the two binding sites strongly suggests an allosteric regulation of the RNAP by (p)ppGpp. Additionally, (p)ppGpp can also bind to other proteins mostly acting as a competitive inhibitor of guanosine nucleotides (9).

A *E. coli* K12 strain devoid of (p)ppGpp synthetase – and thus unable to produce (p)ppGpp hereafter referred as (p)ppGpp^0^ – harbors several physiological defects including amino acids auxotrophy. Growing a (p)ppGpp^0^ strain on minimal medium without amino acids allows the selection of spontaneous suppressors, always mapping to the RNAP, frequently in the β or β’ subunit and rarely in the ω subunit (10). Such RNAP mutants, known as “stringent RNAP” (11), can suppress other (p)ppGpp-related phenotypes.

*Acinetobacter baumannii* is a gram-negative γ-Proteobacterium belonging to the Moraxellaceae family. In 2017, the World Health Organization (WHO) established a list of bacterial pathogens of public health importance and ranked carbapenem-resistant *A. baumannii* (CRAB) amongst the most critical pathogens. More recently in 2024, the list has been revised but CRAB maintained their critical status because of the high mortality following antibiotic-resistant infections as well as the lack of new drug effective against metallo-β-lactamase producing CRAB (12). Known as a multidrug-resistant opportunistic pathogen, *A. baumannii* is also highly resistant and tolerant to stressful environments (13). For instance, clinical isolates of *A. baumannii* can persist for months on dry surfaces in an extreme dehydrated state (14). Intrinsically disordered proteins called hydrophilins were shown to promote tolerance to desiccation (15).

With two long monofunctional RSH enzymes and DksA recently proposed to functionally replace the major stress response sigma factor RpoS (16), *A. baumannii* stands out as unusual and intriguing amongst γ-Proteobacteria. However, studies on (p)ppGpp in *A. baumannii* are still scarce (17,18). As seen in many other pathogens, (p)ppGpp^0^ strains display virulence defects both in mouse and *Galleria mellonella* infection models (17,19). Beyond using serine hydroxamate (SHX) as a proxy for amino acid starvation, the specific stresses that trigger the SR in *A. baumannii* are still unknown. Production of (p)ppGpp has also been described to regulate surface motility with distinct outcomes observed between the reference strains ABUW5075 and ATCC17978 (17,19). This prompted us to further investigate the physiological role of this second messenger in *A. baumannii*.

We were able to identify a third enzyme likely involved in (p)ppGpp metabolism, *ABUW_1957* encoding a SAH capable of hydrolyzing (p)ppGpp *in vivo* both in *E. coli* and *A. baumannii*, although its exact function remains to be elucidated. A classical but thorough phenotyping of RSH mutants highlighted a critical role of (p)ppGpp in the survival of *A. baumannii* cells exposed to nutrient deprivation and desiccation. We also found that iron starvation and exposure to polymyxin B, two stresses possibly encountered during infection, induced (p)ppGpp production. We isolated stringent RNAP mutants that suppress most of the (p)ppGpp-dependent phenotypes, including virulence defects of the (p)ppGpp^0^ strain in the *G. mellonella* model of infection. By combining our suppressor screen with deep sequencing in a Mut-Seq approach (20), we largely expanded our list of candidate missense mutations within *rpoB* and *rpoC* that likely generate stringent RNAP. Finally, we found that ∼9% of the entire transcriptome was significantly modified in (p)ppGpp^0^ cells in contact with solid surface and that the expression of the vast majority of these differentially expressed genes was restored in a stringent RNAP suppressor. Together our data show that RNAP and (p)ppGpp jointly regulate several phenotypical traits critical for the survival in stressful conditions and the virulence of *A. baumannii*.

## Results

### *A. baumannii* harbors three RelA/SpoT homolog enzymes

*A. baumannii* carries two genes coding for long RSH enzymes, *spoT* (*ABUW_0309*; *ABUW_RS01520*) and *relA* (*ABUW_3302*; *ABUW_RS16040*). RelA^Ab^ is a monofunctional synthetase enzyme with a degenerate and inactive hydrolase domain (S^+^/H^-^) (17). However, in contrast to *E. coli* and many other β- and γ-proteobacteria where SpoT is a bifunctional RSH both synthetizing and hydrolyzing (p)ppGpp (S^+^/H^+^), SpoT^Ab^ is a monofunctional hydrolase enzyme with a pseudo-synthetase domain regulating its hydrolase activity (S^-^/H^+^) (4).

In addition to RelA^Ab^ and SpoT^Ab^, we identified two putative short RSH, one SAH (*ABUW_1957*; *ABUW_RS09520*) and one SAS (*ABUW_0769*; *ABUW_RS03770*). First, we investigated the role of the putative SAH hereafter referred to as SahA. In *E. coli*, *spoT* is essential in an otherwise wild-type (WT) background since (p)ppGpp produced by RelA accumulates to a toxic level in a Δ*spoT* background when grown on complex media. In contrast, *spoT* can be inactivated in a Δ*relA* background. To test whether SahA could hydrolyze (p)ppGpp, a P1 lysate made on MG1655 Δ*relA* Δ*spoT::kan^R^* was transduced in a WT strain of *E. coli* MG1655 harboring a pBAD33 plasmid, either empty or expressing *spoT^Ec^*, *spoT^Ab^* or *sahA* under the control of the arabinose-inducible pBAD promoter. As shown in **Fig. S1A**, the endogenous *spoT* gene could be inactivated in the presence of arabinose and one of the three expression vectors (pBAD33-*spoT^Ec^*, pBAD33-*spoT^Ab^* or pBAD33-*sahA*) but not with the empty pBAD33 plasmid, strongly suggesting that SahA is indeed able to hydrolyze (p)ppGpp *in vivo*. Then, we tested whether *sahA* and *spoT* could compensate each other as (p)ppGpp hydrolases in *A. baumannii*. For that, we first generated Δ*relA*, Δ*relA* Δ*spoT*, Δ*relA* Δ*sahA* and Δ*relA* Δ*spoT* Δ*sahA* knock-out mutants in *A. baumannii* in which *relA* is inactivated. Then, a copy of *relA^Ab^* under the control of an IPTG-inducible promoter was introduced in all these mutant strains. Interestingly, induction of *relA^Ab^* in the Δ*relA* Δ*spoT* strain led to a severe growth defect whereas the strain lacking both (p)ppGpp hydrolases (Δ*relA* Δ*spoT* Δ*sahA*) could barely grow when *relA^Ab^* was induced. In contrast, expressing back *relA^Ab^* in strains harboring a functional copy of *spoT –* the single Δ*relA* or the double Δ*relA* Δ*sahA* mutant strain – did not impact the growth. (**Fig. 1**). These results further suggest that SahA is, to some extent, able to hydrolyze (p)ppGpp *in vivo*, both in *E. coli* and *A. baumannii*. Nevertheless, despite several attempts with different methods, we failed to generate a single Δ*spoT* mutant in *A. baumannii*, strongly suggesting that SpoT is the main (p)ppGpp hydrolase in *A. baumannii* and that SahA cannot compensate for the loss of *spoT*, at least in the conditions we tested.

**Figure 1.**
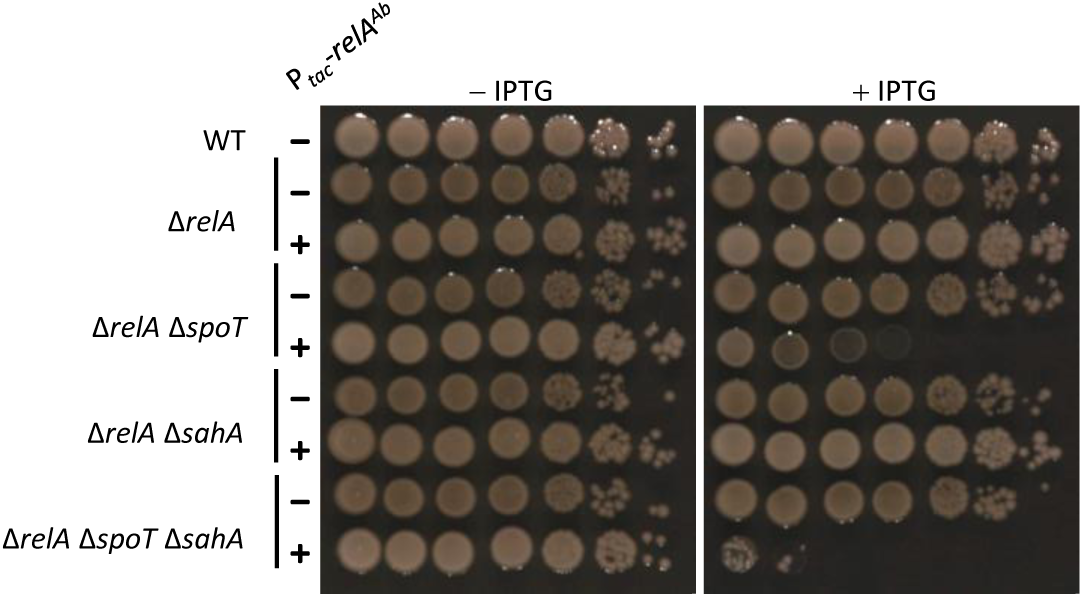
SahA can hydrolyze (p)ppGpp *in vivo* in *A. baumannii.* Viability of *A. baumannii* AB5075 Δ*relA*, Δ*relA* Δ*spoT*, Δ*relA* Δ*sahA,* Δ*relA* Δ*spoT* Δ*sahA* cells carrying a copy of *relA^Ab^* under an IPTG-inducible promoter (P*_tac_*-*relA^Ab^*). Overnight cultures were serial diluted (1:10), spotted on LB agar plates with or without IPTG (500 µM) and incubated overnight at 37 °C.

Finally, we investigated the role of the putative SAS. The fact that a Δ*relA* Δ*spoT* Δ*sahA* triple mutant in *A. baumannii* was viable already suggested that (p)ppGpp was unlikely produced by this putative SAS, at least in our lab conditions. A whole genome sequencing of this triple mutant excluded the presence of any suppressor mutations. To further investigate the potential (p)ppGpp production *in vivo* by this enzyme, *E. coli* MG1655 WT (hydrolase +) and MG1655 Δ*relA* Δ*spoT* (hydrolase -) strains were transformed with a pBAD33 plasmid, either empty or harboring *relA^Ec^* or *ABUW_0769*, and grown on complex LB medium supplemented with glucose or arabinose. Expressing *relA^Ec^* was toxic in the Δ*relA* Δ*spoT* background, likely because of (p)ppGpp accumulation whereas the expression of *ABUW_0769* did not have any impact on the growth of both strains (**Fig. S1B**). Together with the fact that (p)ppGpp was undetectable in the *A. baumannii* Δ*relA* and Δ*relA* Δ*spoT* mutant strains (**Fig. S2)**, these results suggest that no (p)ppGpp was synthesized from ABUW_0769, at least in all the conditions we tested.

### Absence of (p)ppGpp induces pleiotropic effect in *A. baumannii*

It has been previously described that a Δ*relA* mutant in the AB5075 background shows a hyper-motile phenotype and forms elongated cells in stationary phase (17). Considering the high phenotypic heterogeneity among *A. baumannii* strains and also within the same strain in different labs (21) we investigated several known (p)ppGpp-dependent phenotypes in our mutants.

First, the growth of the mutants was measured in complex medium (LB) and minimal synthetic medium with xylose as sole carbon source (MOPSX) and compared to the WT. Surprisingly, we did not find important differences, except a lower plateau for Δ*relA* and Δ*relA* Δ*spoT* cells grown in LB and a slight growth delay in MOPSX (**Fig. 2A and Fig S3A**). Then, we analyzed the cell morphology of bacteria grown overnight in LB. The Δ*relA* and Δ*relA* Δ*spoT* mutants showed heterogenous filamentation with a median of 2.82 and 2.51 μm, respectively, against 2.12 μm for the WT strain (**Fig. 2B and Fig. S3B**). In contrast, the Δ*sahA* mutant showed a cell size distribution close to the WT strain with a median of 2.24 μm (**Fig. S3B**). We also looked at surface motility on low agar plates. After 24 h at 37 °C, the Δ*relA* and Δ*relA* Δ*spoT* strains colonized most of the 90 mm Petri dishes, whereas the WT and Δ*sahA* strains did not move at all, with only growth observed around the inoculation site (**Fig. 2C and Fig. S3C**). Interestingly, the motility behavior correlates well with the colony aspect of the different strains. The WT and Δ*sahA* strains formed white shiny colonies while the Δ*relA* and Δ*relA* Δ*spoT* strains formed browner colonies on LB agar plates (**Fig. S3C**).

**Figure 2.**
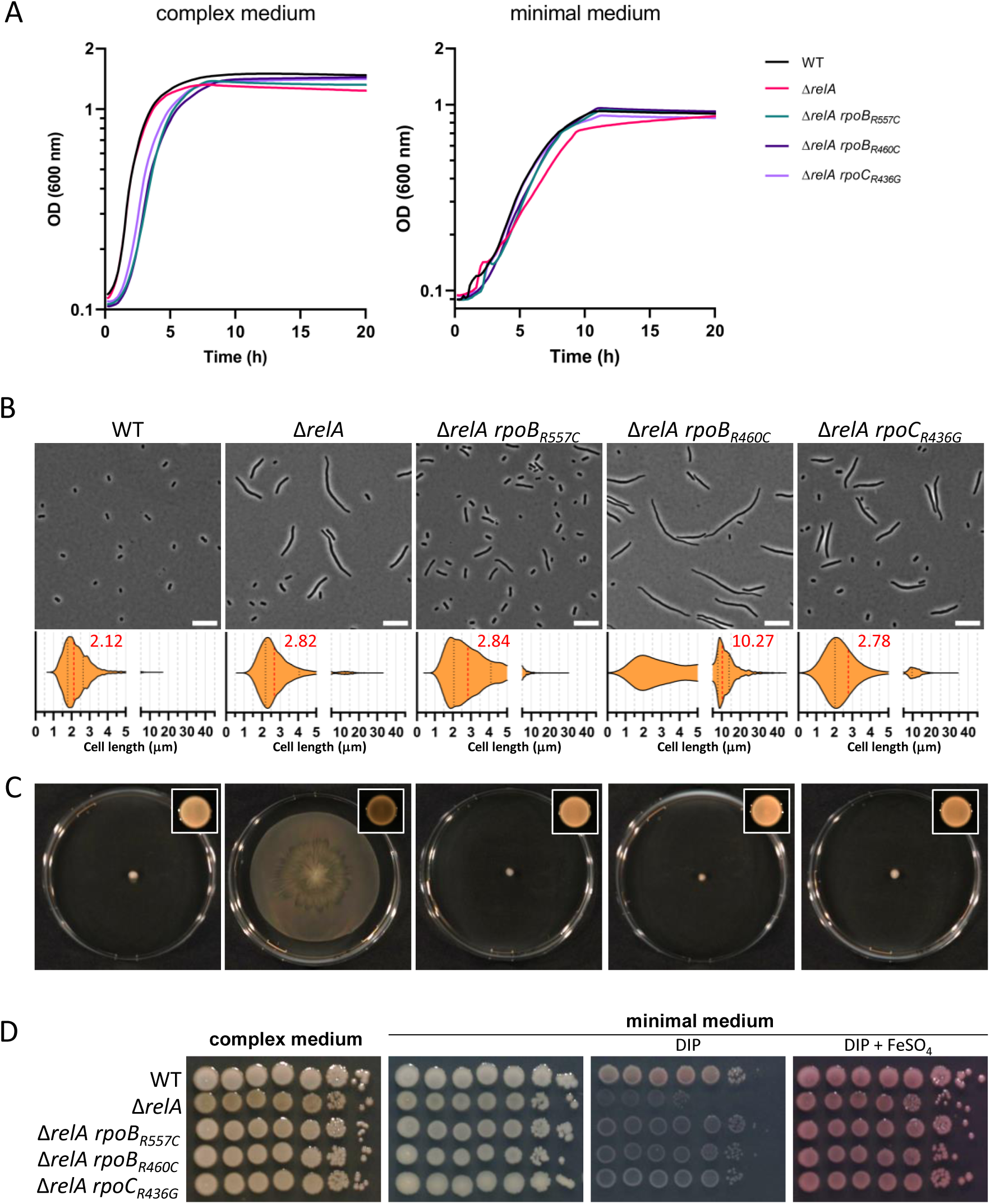
The pleiotropic defects of Δ*relA* are suppressed by mutations in RNAP. (**A**) Representative growth of WT, Δ*relA*, Δ*relA rpoB_R557C_*, Δ*relA rpoB_R460C_* and Δ*relA rpoC_R436G_* at 37 °C in complex (LB) or minimal synthetic (MOPSX) medium for 20 h in 96-well plates. (**B**) Microscopic analysis of WT, Δ*relA*, Δ*relA rpoB_R557C_*, Δ*relA rpoB_R460C_* and Δ*relA rpoC_R436G_* cells grown overnight at 37 °C in liquid complex medium. A representative picture (scale bar = 10 µm) is shown for each strain accompanied by a violin plot showing cell size distribution; the median size (µm) is indicated in red. (**C**) Surface motility of cells described in (B) on low agar (0.5%) complex medium. The appearance and the color of the colonies can also be observed on the picture in the box in top right corners. Pictures for both experiments were taken after 24 h incubation at 37 °C. (**D**) Viability of WT, Δ*relA*, Δ*relA rpoB_R557C_*, Δ*relA rpoB_R460C_* and Δ*relA rpoC_R436G_* strains on complex or minimal synthetic (M9X) medium with or without supplementation [DIP (150 µM); FeSO_4_ (50 µM]. Overnight cultures were serial diluted (1:10), spotted on plates and incubated overnight at 37 °C.

Since *A. baumannii* is characterized by a wide genetic and phenotypic diversity, a Δ*relA* mutant was also constructed in the well-studied ATCC17978 strain. Similarly to AB5075 Δ*relA*, ATCC17978 Δ*relA* formed an heterogenous population of short and longer cells with a median of 2.36 µm when grown overnight in LB compared to 1.75 µm for the ATCC17978 WT strain (**Fig. 3A**). Likewise, we found that in our conditions, WT cells from the ATCC17978 strain were not motile in contrast to the Δ*relA* mutant which was also able to colonize the entire plate in 24 h (**Fig. 3B**).

**Figure 3.**
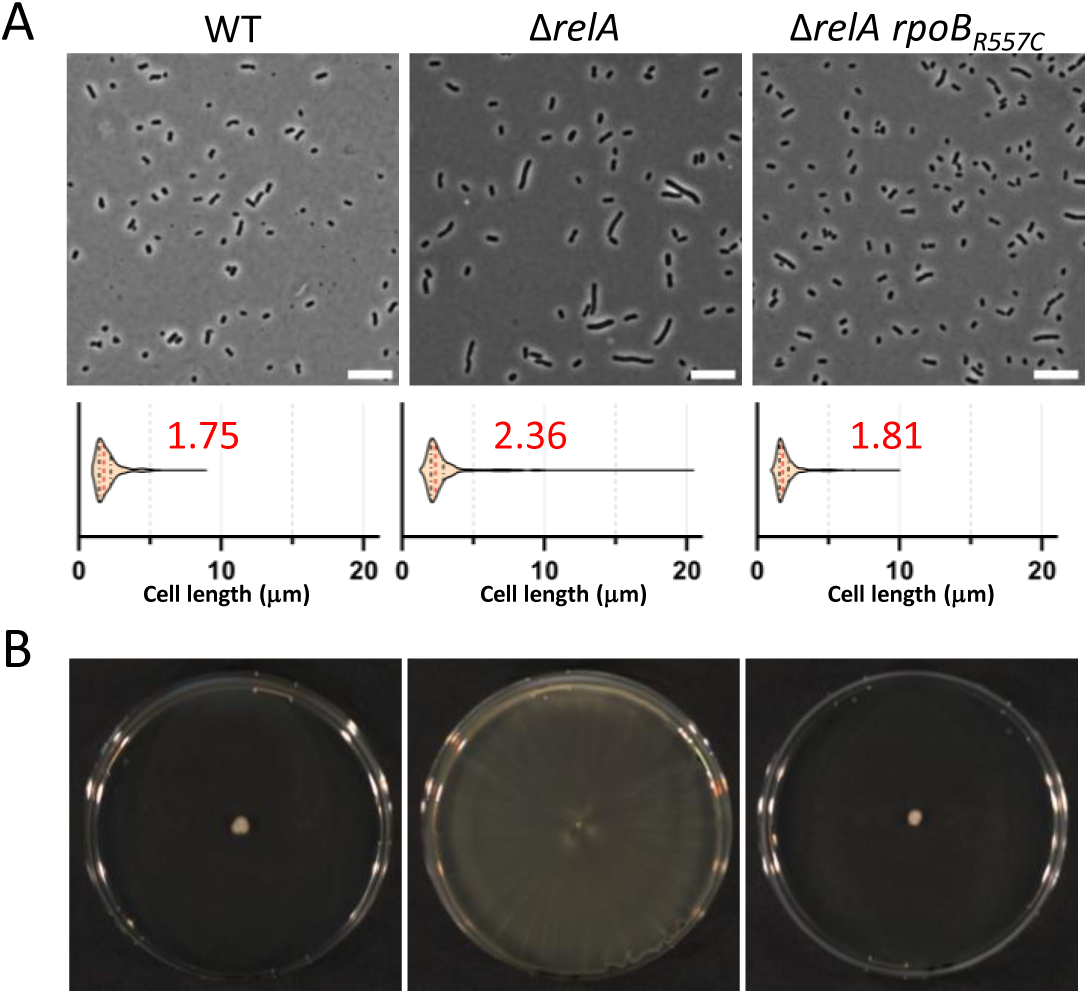
*rpoB_R557C_* is also a suppressor of Δ*relA* in *A. baumannii* ATCC17978 strain. (**A**) Microscopic analysis of WT, Δ*relA*, Δ*relA rpoB*_R557C_ cells grown overnight at 37 °C in liquid LB medium. A representative picture (scale bar = 10 µm) is shown for each strain accompanied by a violin plot showing cell size distribution; the median size (µm) is indicated in red. (**B**) Surface motility of cells described in (A) on low agar (0.5%) complex medium grown at 37 °C for 24 h.

Finally, we used serial dilution drop assays to investigate the sensitivity of our strains to several stresses described as linked to (p)ppGpp. **Figure S3** shows that the Δ*sahA* strain behaved like the WT strain under all conditions tested. Surprisingly, despite the fact that exposure to SHX – known to induce amino acid starvation – strongly induced (p)ppGpp accumulation (**Fig. S2**), the strains unable to produce (p)ppGpp (Δ*relA* and Δ*relA* Δ*spoT*) could grow well on synthetic minimal medium M9 with xylose as sole carbon source (M9X), showing that none of these mutants suffered from amino acid auxotrophy. On the contrary, Δ*relA* mutants had severe growth defects on plates containing triclosan (fatty acid starvation) or 2,2′-dipyridyl (DIP) (iron starvation), while the addition of 50 μM of iron alleviated the defects induced by DIP (**Fig. 2D and Fig. S3C**).

In agreement with our drop assay, we found that DIP led to noticeable ppGpp accumulation. However, to our surprise, neither cerulenin nor triclosan induced (p)ppGpp production in our conditions, suggesting that fatty acid starvation is not a signal triggering (p)ppGpp accumulation (**Fig. 4 and Fig S4**). We also tested polymyxin B and, as previously described for *E. coli* (22), we observed ppGpp accumulation in cells exposed to this last-resort antibiotic used against *A. baumannii* infection (**Fig. 4**).

**Figure 4.**
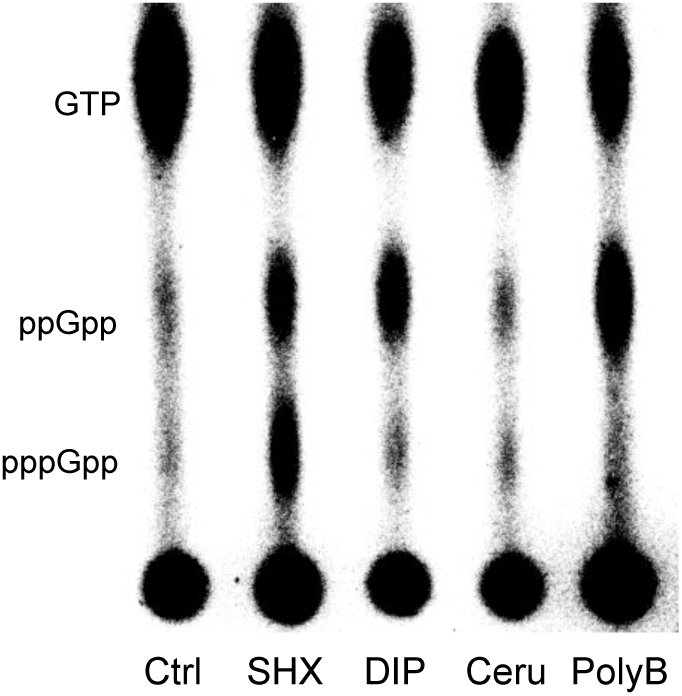
Serine hydroxamate, 2,2’-dipyridyl and polymyxin B induce (p)ppGpp production in *A. baumannii.* Nucleotides extracted from WT cells either non-treated (Ctrl) or exposed to 3.4 mM serine hydroxamate (SHX), 400 µM 2,2’-dipyridyl (DIP), 250 µg/mL cerulenin (Ceru) or 2 mg/mL polymyxin B (PolyB) for 10 min, were analyzed on thin layer chromatography.

### Mutations in *rpoB* or *rpoC* suppress the (p)ppGpp-dependent phenotypes

While characterizing the hyper-motility behaviour of Δ*relA* on low agar plates, we inadvertently selected suppressors, which appeared after more than 96 h of incubation at 37 °C as white shiny dots on top of the bacterial lawn of motile cells (**Figure S5**). Mutations in *rpoB* or *rpoC* – respectively encoding the β and β’ subunit of the RNAP – were found by whole genome sequencing in all tested non-motile suppressor candidates (**Table 1**). In *E. coli*, (p)ppGpp binds to the RNAP to reprogram transcription and mutations in *rpoB* or *rpoC* can induce conformational changes that mimic the effect of (p)ppGpp binding (23). Moreover, some of these so-called stringent mutations were often associated with resistance to rifampicin (Rif^R^). However, none of the suppressor mutations (*rpoB_R460C_*, *rpoB_R557C_* and *rpoC_R436G_*) that we identified here led to a Rif^R^phenotype.

**Table 1.** List of the putative mutations leading to stringent RNAP in *A. baumannii*.

|  | WT AA | Position<br>*A.b* | Position<br>*E.c* | Mutant AA | Hotspot |
| --- | --- | --- | --- | --- | --- |
| RpoB | P | 155 | 153 | L | I |
|  | G | 156 | 154 | V | I |
|  | P | 180 | 178 | L | I |
|  | V | 196 | L194 | G | I |
|  | G | 402 | 401 | V |  |
|  | G | 402 | 401 | R |  |
|  | R | 403 | 402 | P |  |
|  | R | 460 | 451 | H | II |
|  | R | 460 | 451 | C/S | II |
|  | R | 463 | 454 | H | II |
|  | E | 467 | 458 | STOP | II |
|  | S | 521 | 512 | F | II |
|  | A | 541 | 532 | E | II |
|  | L | 542 | 533 | P | II |
|  | G | 543 | 534 | D | II |
|  | G | 543 | 534 | C | II |
|  | G | 545 | 536 | V | II |
|  | G | 546 | 537 | D | II |
|  | A | 552 | 543 | E | II |
|  | R | 557 | 548 | H | II |
|  | R | 557 | 548 | C/S | II |
|  | R | 566 | 557 | P | II |
|  | R | 566 | 557 | G | II |
|  | T | 572 | 563 | I | II |
|  | G | 575 | 566 | C | II |
|  | G | 579 | 570 | S | II |
|  | D | 823 | 814 | E |  |
|  | N | 1093 | 1072 | K |  |
|  | H | 1263 | 1244 | L | III |
|  | T | 1274 | 1255 | R | III |
|  | G | 1280 | 1261 | C | III |
|  | G | 1286 | 1267 | C | III |
|  | Q | 1287 | 1268 | K | III |
|  | L | 1306 | 1287 | F | III |
|  | P | 1336 | 1317 | L | III |
|  | S | 1341 | 1322 | L | III |
|  | S | 1351 | 1332 | P | III |
| RpoC | S | 331 | 326 | F/Y | IV |
|  | R | 342 | 337 | C | IV |
|  | C | 459 | 454 | S | IV |
|  | F | 466 | 461 | C | IV |
|  | A | 499 | 494 | E | IV |
|  | R | 785 | 780 | C | V |
|  | A | 789 | 784 | D | V |
|  | A | 796 | 791 | E | V |
|  | R | 803 | 798 | L | V |
|  | R | 804 | 799 | L/H | V |
|  | A | 1143 | 1147 | D | VI |
|  | R | 1144 | 1148 | C | VI |
|  | V | 1236 | 1240 | G | VI |
|  | G | 1304 | 1308 | D | VI |
|  | R | 1326 | 1330 | C | VI |
|  | G | 1350 | 1354 | D | VI |

Interestingly, these three mutations could suppress most of the (p)ppGpp-dependent phenotypes observed in Δ*relA* (**Figure 2**) except (i) the cell size distribution observed in stationary phase cultures, which was even aggravated for *rpoB_R460C_* (**Fig. 2B**) and (ii) the sensitivity to triclosan, with *rpoC_R436G_* being less resistant than the WT (**Fig. S3D**). More importantly and despite these differences, we found that the three suppressor mutations could fully restore the virulence of *A. baumannii* in the *Galleria mellonella* model of infection. Indeed, the Kaplan-Meier analysis showed that the survival of the larvae infected with each of the suppressors was not significantly different from the WT, while in agreement with previous report (4,17), the larvae infected with Δ*relA* showed a significantly better survival (*p* < 0.0001) (**Figure 5**). These data suggest that the virulence defect of the (p)ppGpp^0^ strain is mainly due to a transcriptional deregulation of virulence factors. In contrast, no significant differences were found when comparing the survival curve of larvae infected with the Δ*sahA* single mutant to the one infected with WT cells (*p* = 0.4022) (**Fig. S6**), suggesting that *sahA* is not required for *A. baumannii* virulence, at least in the *G. mellonella* infection model.

**Figure 5.**
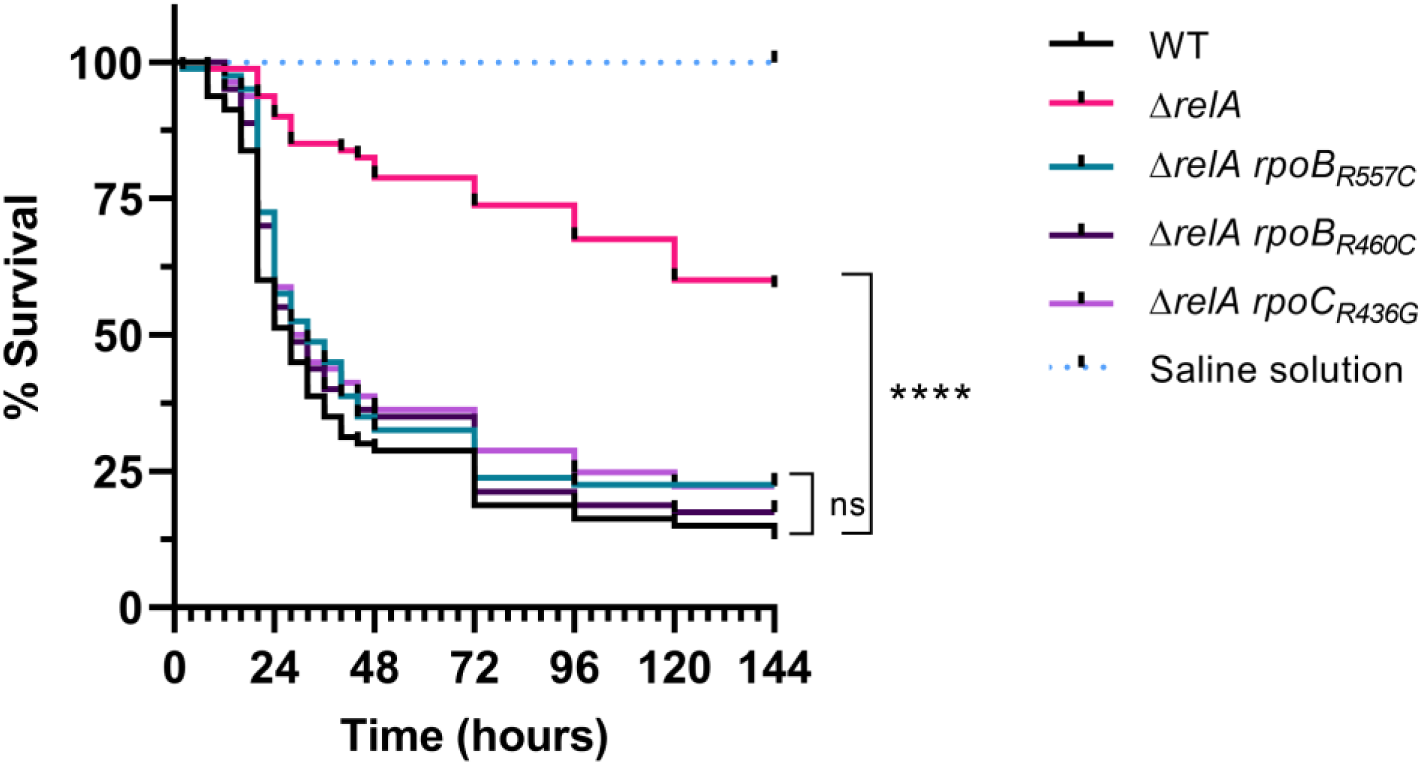
Mutations in *rpoB* and *rpoC* restore the virulence of a Δ*relA* mutant. Kaplan-Meier survival analysis of *Galleria mellonella* larvae infected with the WT, Δ*relA,* Δ*relA rpoB_R557C_*, Δ*relA rpoB_R460C_* and Δ*relA rpoB_R446G_* strains. Each curve represents the average of three independent experiment for a total of 80 larvae per condition. All survival curves are not significantly different from the WT survival curve except the one for the Δ*relA* mutant (Log-rank (Mantel-Cox) test; *p* < 0.0001).

We also selected motility suppressors for the ATCC17978 *ΔrelA* mutants by incubating low agar plates for an extended time, until white shiny colonies appeared on top of the bacterial lawn as observed for the AB5075 strain. By using specific primers that anneal to the *rpoB_R557C_* allele, we were able to fish out 8 positive clones that were then validated by Sanger sequencing. Again, as for AB5075, *rpoB_R557C_* was able to suppress most of the phenotypes displayed by Δ*relA* in the ATCC17978 strain (**Fig. 3**).

To check to what extent *rpoB* and *rpoC* are hotspots for the selection of Δ*relA* suppressors, we used an unbiased Mut-Seq approach (20) to map as many mutations as possible in *rpoB* or *rpoC* that suppress the hyper-motility phenotype displayed by Δ*relA*. For that, 284 non-motile Δ*relA* suppressors were selected on low agar complex medium plates as described above. It is important to note that among them, only 8 were Rif^R^ (**Table 1**). Then, the pooled gDNA of these 284 clones was used as a template to amplify *rpoB* and *rpoC* loci and both PCR products were deep sequenced. By using a confidence threshold of 0.5% (see M&M section), 39 and 18 putative substitutions – leading to 32 and 16 missense mutations respectively in *rpoB* and *rpoC* – were identified (**Fig. 6 and Table 1**).

**Figure 6.**
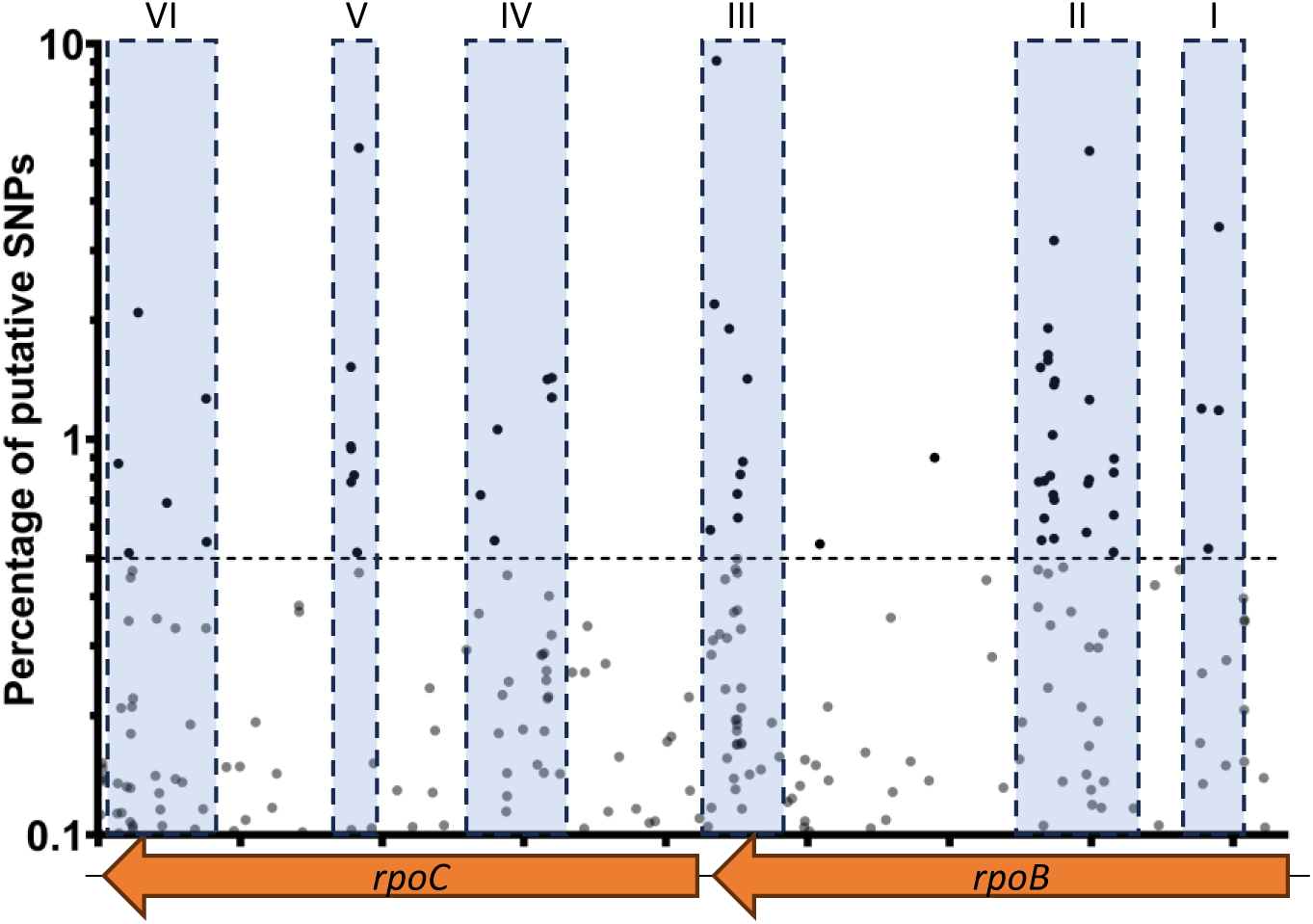
Putative stringent mutations leading to loss-of-motility in Δ*relA* are spread in 6 sub-regions within *rpoB* and *rpoC.* For a given nucleotide position, only SNPs found by Mut-Seq with a percentage ≥ 0.1 are represented. Confidence threshold of 0.5% is represented by the horizontal dashed line. Blue boxes highlight the sub-regions where most SNPs were found.

Together these data strongly support that the (p)ppGpp-dependent transcriptional reprogramming is responsible for most of the phenotypes observed in (p)ppGpp^0^ cells and suggest that RNAP is a primary (p)ppGpp target in *A. baumannii*.

### A single substitution in *rpoB* can revert the (p)ppGpp-dependent transcriptomic changes

To further characterize the suppressing effects of *rpoB_R557C_* in Δ*relA*, the RNA extracted from the WT, Δ*relA* and Δ*relA rpoB_R557C_* cells grown on the medium used for the selection of the suppressors – LB low agar plates – was sequenced. When comparing Δ*relA* to the WT, we found 304 (8.09%) differentially expressed genes (DEGs) (absolute log_2_ Fold Change |log_2_FC| ≥ 2 and FDR–adjusted *p*-value ≤ 0.005), with 115 (3.06%) up-regulated genes and 189 (5.03%) down-regulated genes (**Table S1**). As expected, the *ABUW_3766-3773* operon known to be critical for surface motility (24,25) was found among the most up-regulated genes in the Δ*relA* strain (**Fig. 7A**). Next to this operon, the *abaRMI* (*ABUW_3774-3776*) quorum-sensing locus was also highly up-regulated. Genes related to iron homeostasis (acinetobactin gene cluster and several TonB-dependent siderophore receptor) and fatty acid (FA) biosynthesis were also up-regulated in the Δ*relA* mutant (**Fig. 7A** and **Table S1**). Several genes known to be regulated by (p)ppGpp in *E. coli* (*ostAB*, *raiA* and *fis*) were found to be down-regulated in Δ*relA*. In addition, the expression of the locus *ABUW_2433-2443*, containing stress related genes such as *cinA1*, *katE, dtpA and dtpB* was also found to be down-regulated in Δ*relA*. Apart from these, most of the other down-regulated genes were of unknown functions (**Fig. 7A** and **Table S1).**

**Figure 7.**
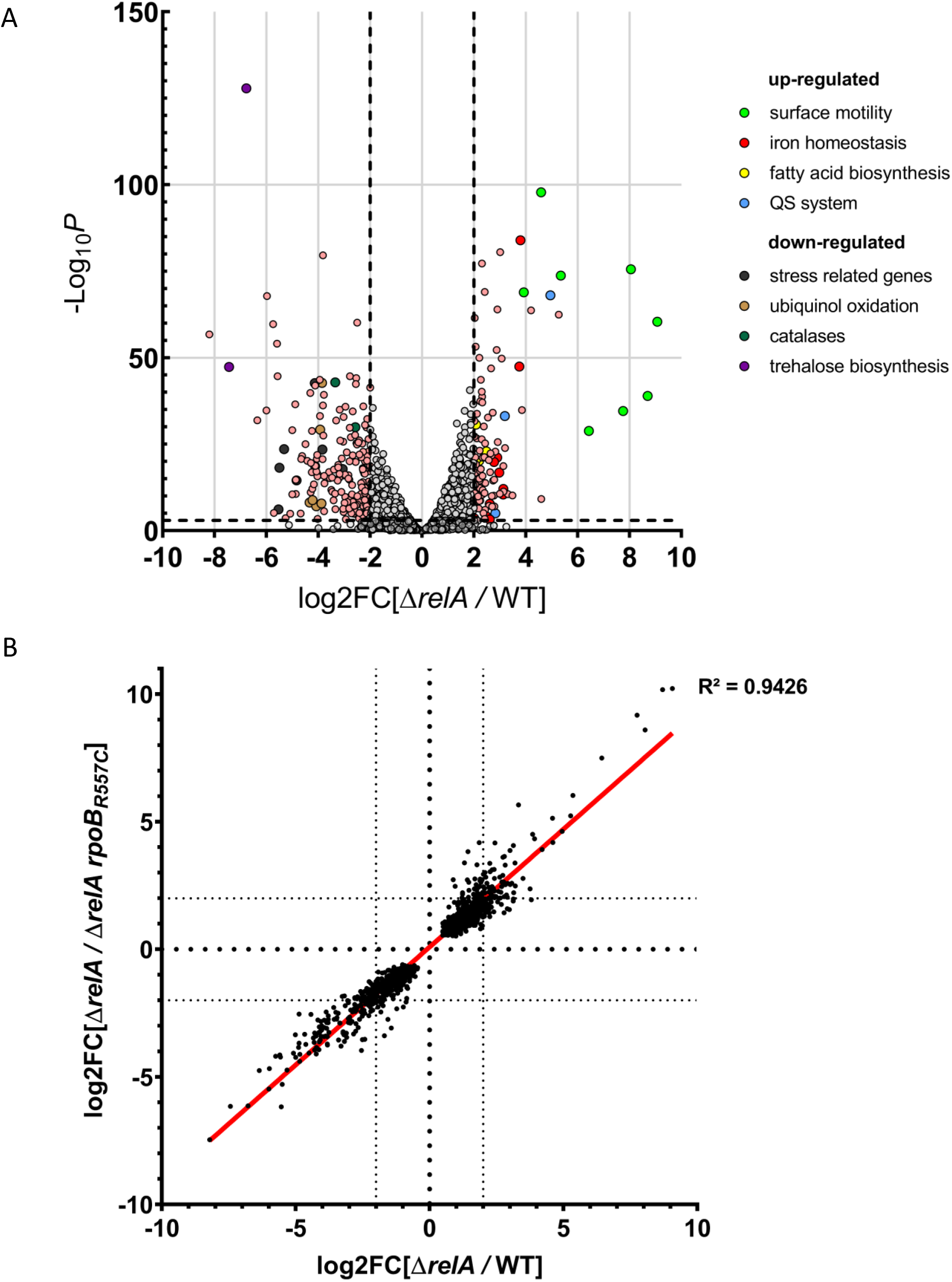
The *rpoB_R557C_* mutation reverts back the transcriptomic profile of the Δ*relA* mutant close to a WT on low agar complex medium. **(A)** Volcano plot representing the genes differentially expressed in the Δ*relA* mutant compared to the WT strain on low agar medium. Genes with a |log_2_FC| ≥ 2 and –Log_10_ *p*-value ≥ 3 are represented in light pink or with a color code for group of genes associated with known functions (light green: surface motility; red: iron homeostasis; yellow: fatty acid biosynthesis; blue: quorum sensing (QS) system; black: stress related genes; brown: ubiquinol oxidation; purple: trehalose biosynthesis; dark green: catalases). **(B)** xy-plot representing the differential expression of genes between the Δ*relA*/WT *vs* Δ*relA/*Δ*relA rpoB_R557C_* on motility complex medium. Each dot represents the log_2_FC value for the strains indicated under brackets (only genes with FDR-adjusted *p* ≤ 0.005 were kept).

Interestingly, we only found 21 (0.56%) DEGs between the WT and the suppressor Δ*relA rpoB_R557C_* strain, with respectively 15 (0.40%) up- and 6 (0.16%) down-regulated genes. **Figure 7B** shows that the log_2_FC values of Δ*relA vs* WT and Δ*relA vs* Δ*relA rpoB_R557C_* have a linear correlation (R² = 0.9426). This indicates that the transcriptomic changes induced in the Δ*relA* mutant are almost entirely reverted to WT in the Δ*relA rpoB_R557C_* suppressor.

We also determined the (p)ppGpp-dependent transcriptome for cells grown in liquid LB medium by comparing WT to Δ*relA* or Δ*relA rpoB_R557C_* strains. However, in these conditions only 153 genes (4.06%) were differentially expressed in Δ*relA* compared to WT (absolute log_2_ Fold Change |log_2_FC| ≥ 2 and FDR–adjusted *p*-value ≤ 0.005), with 28 (0.74%) up-regulated genes and 125 (3.32%) down-regulated genes (**Fig. S7A** and **Table S2**). In particular all the genes encoding the assembly machinery of Csu pili (*csu* genes) were down-regulated in Δ*relA* (**Fig. S7A**). Importantly, there were still 165 DEGs when comparing the suppressor Δ*relA rpoB_R557C_* to the WT strain (**Table S2)**, suggesting that the *rpoB_R557C_* restored the (p)ppGpp-dependent transcriptional profile more efficiently on low agar plate than in liquid medium. In agreement with this, the linear regression between the log_2_FC values of Δ*relA* and WT and Δ*relA* vs Δ*relA rpoB_R557C_* did not fit well for the liquid medium (R² = 0.4531; **Fig. S7B**), showing a low correlation. These data suggest that the contact of *A. baumannii* onto a semi-solid surface could induce (p)ppGpp accumulation, which leads to a deep transcriptional reprogramming.

### The Δ*relA* mutant is highly sensitive to desiccation and have impaired H_2_O_2_ detoxification

Among the genes differentially expressed in the Δ*relA* mutant in both liquid and low agar media, we identified the TetR-type transcriptional regulator *ABUW_1645*. It was the most down-regulated gene in liquid medium (log_2_FC = -7.91) and among the most down-regulated on low agar plates (log_2_FC = -5.27). This regulator has been previously described as being involved in the pleiotropic phenotypic switch in *A. baumannii* (26). Since its expression appears to be partially restored in the Δ*relA rpoB_R557C_* suppressor strain compared to the WT (log_2_FC of -0.61 in liquid and -2.32 in low agar medium), we wondered whether it could contribute to phenotypes observed in the Δ*relA* mutant. To test this hypothesis, a second copy of *ABUW_1645* expressed from the IPTG-inducibe P*_tac_* promoter (P*_tac_*-*ABUW_1645*) was introduced in the Δ*relA* mutant as well as in the WT. However, this construct turned out to be lethal on IPTG containing plates, not only in the Δ*relA* mutant but also in the WT strain, while it did not impact the growth or the colony appearance without IPTG. Occasionally, WT and Δ*relA* clones with P*_tac_*-*ABUW_1645* were able to grow in the presence of IPTG, but these clones harbored IS4 ISAba1 transposase inserted into *ABUW_1645*, thereby disrupting its function. Although we cannot firmly exclude the possibility that *ABUW_1645* participates to some extent to the phenotypes displayed by Δ*relA*, our data show that *ABUW_1645* is toxic when highly expressed.

On low agar medium, the most down-regulated gene in the Δ*relA* strain is *dtpA* (log_2_FC = -8.21), which encodes a hydrophilin involved in the tolerance of the *Acinetobacter calcoaceticus–baumannii* complex to high desiccation (15). Since the *dtpA* locus (*ABUW_2433–2443*) is down-regulated in both liquid and low agar media, and includes several stress-related genes such as those encoding the second hydrophilin DtpB and the catalase KatE, we measured the tolerance to desiccation and the catalase activity in the WT, Δ*relA* and suppressors strains. The Δ*relA* strain showed a significant decrease (*p* < 0.0001) in normalized catalase activity (37.4% ± 10.6%) compared to the WT strain (100% ± 8.0%). All the three Δ*relA* suppressors displayed a normalized catalase activity close to or above the WT, with 138.4% ± 11.5% (*rpoB_R557C_*), 97.0% ± 9.1% (*rpoB_R460C_*) and 122.2% ± 9.3% (*rpoC_R436G_*) (**Fig. 8A**). We then tested the survival of the same strains upon desiccation. After three days at 35% room humidity, 4.5% ± 2.5% of CFU were recovered for the WT strain while only 0.009% ± 0.005% CFU were recovered for the Δ*relA* mutant. Compared to the Δ*relA* mutant, all the three suppressors showed increased desiccation survival although with some differences between them in the CFU recovering (3.8% ± 3.7% for *rpoB_R557C_*, 2.4% ± 0.6% for *rpoB_R460C_* and 0.12% ± 0.08% for *rpoC_R436G_*) (**Fig. 8B**).

**Figure 8.**
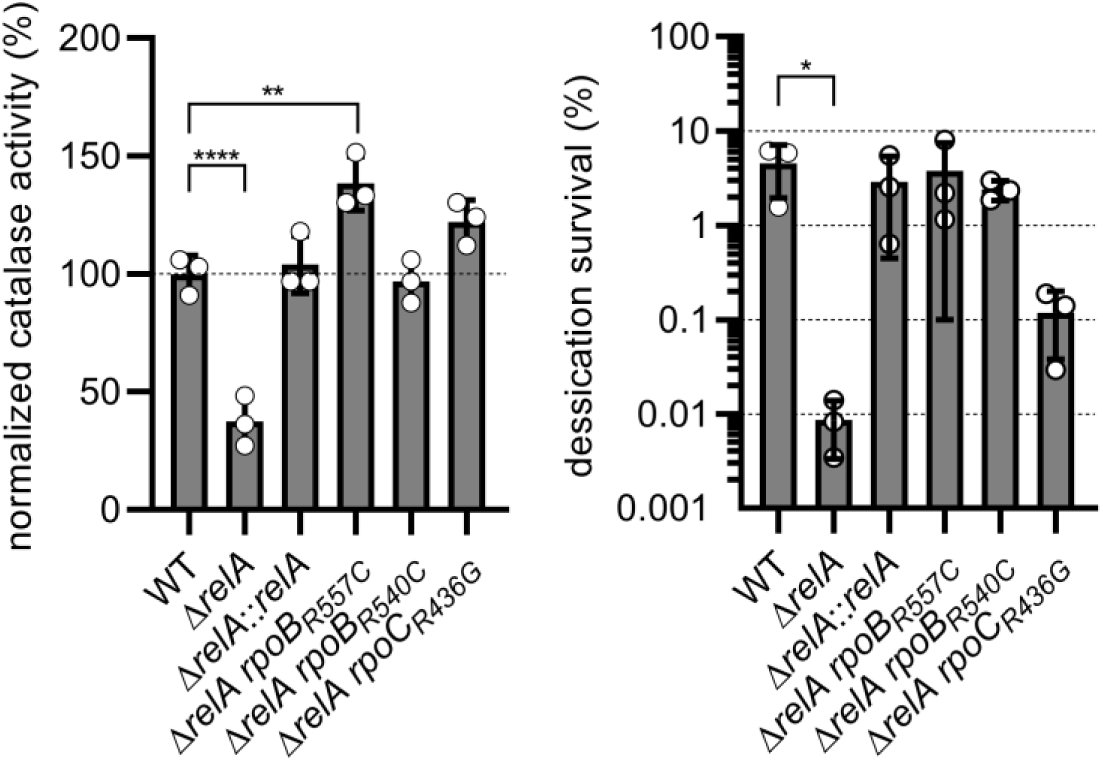
The Δ*relA* mutant has impaired catalase activity and desiccation resistance. **(A)** Catalase activity produced by overnight culture of the WT, Δ*relA,* Δ*relA*::*relA*, Δ*relA rpoB_R557C_*, Δ*relA rpoB_R460C_* and Δ*relA rpoB_R436G_* strains. ANOVA and Dunnet’s multiple comparisons test (p < 0.01 and *p* < 0.0001). **(B)** Desiccation survival of the same strains after 3 days at 35% room humidity and 25 °C. Kruskal-Wallis and Dunn’s multiple comparisons test (*p* < 0.05).

## Discussion

In this study, we investigated and characterized the pleiotropic roles played by (p)ppGpp in *A. baumannii*. First, we identified potential new RSH enzymes including a putative orphan SAS (*ABUW_0769*) that might synthesize (p)ppGpp. However, we did not find any correlation between its expression and (p)ppGpp production neither in *E. coli* nor in *A. baumannii*. Nevertheless, we cannot rule out the possibility that a specific condition is required to activate (p)ppGpp synthesis by ABUW_0769. We also identified a putative orphan gene (*ABUW_1957*) coding for a SAH (SahA) that was able to hydrolyze (p)ppGpp both in *E. coli* and *A. baumannii*. However, under the tested conditions, its hydrolase activity alone was insufficient to counterbalance the absence of *spoT* and to counteract (p)ppGpp produced by RelA. During our phenotypic analysis of *A. baumannii* RSH enzymes, we did not identify a condition where the Δ*sahA* mutant behaved differently from the WT strain, including virulence in *G. mellonella*. Small alarmone hydrolases remain poorly characterized in bacteria but recently, two orphan SAHs have been described in Proteobacteria. In addition to (p)ppGpp, SAHs were shown to degrade another alarmone, (p)ppApp, and were suggested to act as a defensive mechanism during bacterial competition (27–29). SahA could potentially play a similar role during host colonization, but the involvement of small RSHs in bacterial competition is still an emerging field, requiring further investigation.

Then, we extended our analysis to the long RSHs, RelA and SpoT. All attempts to generate a single *spoT* mutant failed, suggesting that SpoT is essential in *A. baumannii*, even in the presence of SahA. In all β- and γ-Proteobacteria studied so far, differences have been observed between a single Δ*relA* mutant and a double Δ*relA* Δ*spoT* mutant, essentially because SpoT is bifunctional. However, in *A. baumannii*, both RSHs are monofunctional, with RelA working as the sole (p)ppGpp synthetase and SpoT as a hydrolase (4). We found that both mutants exhibited the same phenotypes in all the tested conditions, both behaving as (p)ppGpp^0^ strains as expected. Therefore, our results exclude any (p)ppGpp-independent role played by the SpoT protein in *A. baumannii*, at least in all our conditions.

The absence of (p)ppGpp activated surface motility on low agar plates and altered colony morphology, likely reflecting a change of the cell surface properties. The lack of (p)ppGpp also had a detrimental effect on cellular morphology, leading to filamentation of a part of the population in stationary phase. Finally, as already observed with all pathogens studied so far, the virulence of *A. baumannii* in *G. mellonella* was also impacted in the (p)ppGpp^0^ strains. To expand our analysis, we also generated a (p)ppGpp^0^ strain in *A. baumannii* ATCC17978 and we obtained similar results in both AB5075 and ATCC17978 backgrounds, including the surface motility since both WT were not motile while (p)ppGpp^0^ mutants were hyper-motile. This was surprising considering that *A. baumannii* ATCC17978 WT strain was often reported in the literature as motile and a Δ*relA* mutant has been shown in one report to be less motile than the WT strain (19). Aside from lab conditions and experimental setup, these differences could also be explained by the fact that there are at least two main laboratory variants of the *A. baumannii* ATCC17978 strain (21), the primary difference being the presence or absence of a 44-kb locus (AbaAL44) known to be responsible for several phenotypic changes including surface motility (21). In addition to the AbaAL44 locus, several SNPs have been also identified, including a mutation in *obgE*, a GTPase that may be involved in the SR and thus have an impact on surface motility (30).

While challenging the (p)ppGpp^0^ strains with stressful conditions known to be (p)ppGpp-dependent in other species, we found that in the absence of (p)ppGpp, *A. baumannii* remained prototrophic for amino acids when grown in synthetic minimal medium with xylose as sole carbon source. However, (p)ppGpp strongly accumulated upon SHX treatment, indicating that the hallmark of the SR is functional in *A. baumannii*. We also observed that FA and iron starvation both affected the growth of (p)ppGpp^0^ cells whereas only iron limitation led to (p)ppGpp accumulation. In *E. coli*, FA starvation leads first to a SpoT-dependent (p)ppGpp synthesis (31). Given that the SpoT SD domain is degenerated and non-functional in *A. baumannii*, this could explain the lack of (p)ppGpp production upon triclosan exposure. In *E. coli*, a prolonged FA starvation ultimately leads to the depletion of pyruvate used as a precursor for lysine synthesis, which thereby triggers the (p)ppGpp synthesis by RelA (32). However, this regulation has only been described in *E. coli* and may differ in *A. baumannii*. Notwithstanding the absence of (p)ppGpp accumulation upon FA starvation, our transcriptomic analyses show that genes coding enzymes involved in FA biosynthesis are upregulated in Δ*relA*, suggesting that (p)ppGpp^0^ cells behave like being starved for FA even in rich media. At least these data exclude the possibility that the growth defects of Δ*relA* cells in the presence of triclosan comes from low levels of FA biosynthetic enzymes.

Iron deprivation also leads to (p)ppGpp accumulation in *E. coli* but in contrast to FA starvation, only SpoT is involved with its activity switching toward (p)ppGpp synthesis over hydrolysis (33). In *A. baumannii* however, (p)ppGpp production during iron starvation can only result from (p)ppGpp synthesis by RelA. Two not mutually exclusive scenarios are therefore possible. Either the SpoT-dependent regulation is conserved, which leads to a decrease of (p)ppGpp hydrolysis and a concomitant increase of (p)ppGpp levels due to basal synthetase activity of RelA, which cannot be hydrolyzed. Or iron starvation indirectly leads to amino acid(s) starvation, which in turn activates (p)ppGpp synthesis by RelA.

During the characterization of surface motility, we observed an intriguing phenomenon since after prolonged incubation, white and shiny colonies appeared on the bacterial motility lawn of (p)ppGpp^0^ cells, strikingly resembling to WT colonies. These extragenic suppressors are RNAP mutants that not only reverted surface motility but also suppressed most phenotypes associated with the loss of (p)ppGpp, including virulence in *G. mellonella*. Surprisingly, the *rpoB_R460C_* mutation was as efficient as *rpoB_R557C_* and *rpoC_R436G_* to suppress the lack of virulence of Δ*relA* despite the very strong filamentation observed in stationary phase. Of course, we cannot rule out the possibility that this filamentation was not induced inside *G. mellonella*.

We applied our motility suppressor screen combined with high-throughput sequencing (Mut-Seq) to isolate as many mutations as possible in RNAP. With this approach, 48 putative missense mutations were identified in *rpoB* or *rpoC*, among which we found the three mutations we had previously isolated (*rpoB_R460C_*, *rpoB_R557C_* and *rpoC_R436G_*). In 2006, Trinh and colleagues undertook the enormous task of compiling most RNAP mutations reported so far across various organisms, including *E. coli* (23). Among these were also mutations mimicking (p)ppGpp effect, known as “RNAP stringent mutations”. We compared our list of mutations obtained with Mut-Seq to this reference list and found that more than 50% of the stringent mutations described in *E. coli* were also present under our conditions in *A. baumannii*, strongly suggesting that our screen was effective in selecting for stringent mutations, including putative new mutations that we believe have never been described before.

With a total of 57 different substitutions identified within *rpoB* and *rpoC* in our screen, and in agreement with work done on *E. coli*, RNAP seems to be the primary target for mutations in the (p)ppGpp^0^ background in *A. baumannii*. In support of that, by comparing the WT, Δ*relA*, and Δ*relA rpoB_R557C_* strains on surface motility medium, we found that (i) the expression of many genes (∼8%) is regulated by (p)ppGpp and (ii) their expression was mostly reverted close to the WT level with the *rpoB_R557C_* mutation. In contrast, the (p)ppGpp-dependent regulon was much smaller (∼4%) when strains were sampled from mid-exponential phase of growth in liquid complex medium and *rpoB_R557C_* poorly suppressed these transcriptional changes. Together, these data suggest that the low agar medium used for surface motility is “stressful” for *A. baumannii*, at least more than the liquid complex medium. In support of this, the *ABUW_2433-2444* locus containing stress-related genes linked to oxidative stress and desiccation was highly up-regulated in the WT strain (mean log_2_FC locus = 5.34 ± 2.11) grown on semi-solid medium compared to liquid medium. On the contrary, the expression of the hydrophilins (*dtpA* and *dtpB*) and the catalase (*katE*) encoded in this locus was not induced in (p)ppGpp^0^ cells, which led to a poor desiccation resistance and catalase activity. Strikingly, the suppressor mutations (*rpoB_R460C_*, *rpoB_R557C_* and *rpoC_R436G_*) were all able to suppress these phenotypes, which corroborates the high expression of the *ABUW_2433-2444* locus observed in the Δ*relA rpoB_R557C_* strain.

Altogether our data highlight a key role played by the second messenger molecule (p)ppGpp in the virulence and survival upon stress of the opportunistic pathogen *A. baumannii*.

## Materials and Methods

### Bacterial strains and growth conditions

Strains, plasmids and oligonucleotides used in this study are listed in **Table S3**. Details for plasmids construction are presented in supplementary methods. *E. coli* and *A. baumannii* strains were grown at 37 °C either aerobically with shaking at 175 rpm or on plate with 1.5% agar (Difco™). LB Broth Base (Invitrogen™) medium was used as complex medium for both *E. coli* and *A. baumannii*. M9 and MOPS were used as a synthetic minimal medium with xylose 0.3% (M9X and MOPSX, respectively) as carbon a source for *A. baumannii*. M9 was prepared using M9 salts (49 mM Na_2_HPO_4_, 22 mM KH_2_PO_4_, 8.6 mM NH_4_Cl 18.6 mM NH_4_Cl) supplemented with 1 mM MgSO_4_ and 0.1 mM CaCl_2_. When needed 0.5% glucose, 0.5% arabinose or 500mM/1mM IPTG were added.

Growth was monitored by measuring absorbance at OD_600 nm_ in liquid cultures using an automated plate reader (Biotek, Epoch 2) with continuous linear shaking (700 rpm) for at least 20h at 37 °C.

For *E. coli*, antibiotics were used at the following concentrations (μg/mL; in liquid/solid medium): chloramphenicol (30/30), apramycin (30/30) while for *A. baumannii*, media were supplemented with apramycin (30/30) or rifampicin (20/20) when appropriate. Plasmid delivery into *A. baumannii* was achieved by natural transformation as described in (34). In-frame deletions were created by using the pK18-*mobsacB* derivative plasmids as follows. Integration of the plasmids in the *A. baumannii* genome after single homologous recombination were selected on LB plates supplemented with apramycin. Independent recombinant clones were then inoculated in LB medium without antibiotics and incubated overnight (ON) at 37 °C. Then, dilutions were spread on LB agar plates without NaCl and supplemented with 10% sucrose and incubated at 30 °C. Single colonies were picked and transferred onto LB agar plates with and without apramycin. Finally, to discriminate between mutated and wild-type loci, apramycin-sensitive clones were tested by PCR on colony using locus-specific oligonucleotides.

### Microscopy experiment

Bacteria were grown ON directly inoculated from -80 °C stocks. Five microliters of ON culture diluted to ½ were spotted on a LB agar pad. Phase contrast images were obtained using Axio Observer 7 microscope (ZEISS, Germany), Orca-Flash 4.0 camera (Hamamatsu, Japan) and Zen 3.9 software (ZEISS, Germany). Analysis was done with the MicrobeJ plugin (35) on Fiji software (36) in order to determine the cell length.

### Motility assay

Motility medium containing LB supplemented with 0.5% agar (Difco™) was prepared the day prior of the experiment and plates were poured extemporaneously. A single isolated colony was used to inoculate a single plate with a sterile toothpick. Plates were incubated during 24 h at 37 °C.

### Measurement of (p)ppGpp

Bacteria were grown ON directly inoculated from -80 °C stocks in MOPSX and diluted in MOPSX low phosphate to OD ∼ 0.1 and grown until OD = 0.4-0.6 and then diluted again to OD 0.1 in the same medium. Then radiolabeled phosphate [γ32P]-KH_2_PO_4_ was added (final concentration: 40 µCi) and cells were incubated at 37 °C - 600 rpm. After 15 min, stressors (final concentrations: 3.4 mM serine hydroxamate, 400 µM 2,2’-dipyridyl, 250 µg/mL cerulenin, 250 µg/mL triclosan and 2 mg/mL polymyxin B), their solvent alone (final concentration: 2% ethanol) or MOPSX low phosphate was added and cells were incubated again during 10 to 30 min. To extract nucleotides, 200 µL of cells were added to 80 µL ice cold 50% formic acid and kept and ice for 20 min. Then, tubes were kept at -20 °C during at least 1 h. Nucleotides were migrated on polyethyleneimine (PEI) cellulose plate (Macherey-Nagel, Duren, Germany) in 1.5 M KH_2_PO_4_ (pH 3.4) at room temperature. TLC plates were air dried and exposed against MS Storage Phosphor Screen (GE Healthcare) overnight. Phosphor screens were finally visualized using an Amersham Typhoon (Cytiva).

### Virulence Assay

The animal model used in this study is based on the larvae of the insect *Galleria mellonella.* Bacterial strains of *A. baumannii* were cultured ON in LB broth at 37 °C, harvested 5 min at 4500 rpm and washed twice in saline solution (0.9% NaCl). Cultures were adjusted to reach a density of ∼10^6^ CFU/larva and injected into the last proleg of *G. mellonella* larvae of ∼0.215 g each, using a 26 G needle (Terumo, Belgium) and a syringe pump (Thermofisher, MA, USA). The bacterial inoculum size was verified by plate counting on LB agar. Larvae were housed five per Petri dish and incubated at 37 °C for 6 days. Live and dead larvae were counted every 4 h for 48 h post-infection and then once a day for up to 6 days. Three independent experiments were performed to treat a total of 80 animals per condition. Data are presented on Kaplan-Meier curves, and log-rank statistical analysis was performed with GraphPad Prism 9.0 software.

### Suppressors and Mut-Seq

Suppressors were selected on surface motility (LB low agar) plates by incubating at least 96 h at 37 °C. Then single colonies were isolated, incubated in LB, grown ON at 37 °C. ON cultures were used to extract genomic DNA (NucleoSpin Tissue, Macherey-Nagel, Duren, Germany). For the Mut-Seq, six isolated colonies were used to inoculate 30 surface motility plates. Ten putative suppressors were picked on each plate and inoculated in 96 well plates. Each well was tested for surface motility, growth on LB agar plate with or without rifampicin. Overnight cultures of ∼300 putative suppressors were done in 96 well plates, then 10 µL culture for each clone was pooled and used to extract genomic DNA. The *rpoB* and *rpoC* genes were amplified with pair of primers 4152/4153 and 4154/4155 respectively, on the genomic DNA mix, with Q5^®^ High-Fidelity DNA polymerase (New England BioLab) according manufacturer recommendation. Both templates were mixed at a 1:1 ratio. Both Whole Genome Sequencing (suppressors) and DNA sequencing (Mut-Seq) were performed using an Illumina NextSeq 2000 (paired-end 2 x 100) instrument (Seqalis, Gosselies, Belgium). Data analysis was performed using HISAT2 and Rsamtools on Galaxy and R software, respectively.

### Transcriptomic analysis

Total RNAs were extracted either from growing bacteria in liquid culture at comparable cell density (OD_600 nm_ ∼ 0.5) or from bacterial colonies on motility medium. At least two independent biological replicates were made for each strain. Extraction was performed using RNeasy mini kit (QIAGEN) according to the manufacturer’s instructions. RNASeq TTRNA libraries were prepared and sequenced with Illumina NextSeq 2000 (paired-end 2x100) instrument (Seqalis, Gosselies, Belgium). NGS data were analysed using Galaxy (https://usegalaxy.org) (37) as described in (38). Briefly, FastQC/Falco was used to evaluate the quality of the reads; HISAT2 was used to map the reads onto the AB5075 reference genome (NZ_CP008706) and generate bam files; featureCounts was used to generate counts tables using bam files and DESeq2 was used to determine differentially expressed genes. The Volcano plot was generated using GraphPad Prism 9 software.

### Catalase Assay

Catalase assay was performed as described in (39). Briefly, bacteria were grown ON directly inoculated from -80 °C stocks. Cells were then washed and normalized to OD ∼ 4. In a glass tube (15 mm diameter x 130 mm height), 100 µL of cells, 100 µL of 1% Triton X100 and 100 µL of undiluted hydrogen peroxide (27%) were mixed. The heigh of the foam was measured after 5 min and subtracted by the height of the control without cells.

### Desiccation Assay

Bacteria were grown ON directly inoculated from -80 °C stocks. On a 24-well plate, for each strain, 25 µL of cells was deposited in 3 different wells in a randomized manner and allowed to dry during 30 min in a microbiological safety cabinet and then incubated at 25 °C with 35% room humidity. Cells were harvested either directly after drying for the inoculum or after 3 days. To harvest cells, 300 µL of M9 salt was added in each well during 30 min to allow cell rehydration. Then cells were diluted and plated on LB medium to determine the number of colony forming units.

### Statistical analyses

All the statistical analyses were performed using GraphPad Prism 9 software.

### Data availability

RNA-Seq and Mut-Seq data have been deposited to the Gene Expression Omnibus (GEO) repository.

## Acknowledgements

We thank Charles Van der Henst for supplying *A. baumannii* WT strains and Paul Guiraud for help with the microscopy experiments. This work was supported by a Welbio Starting Grant (WELBIO-CR-2019S-05) to R.H. A.P. was a postdoctoral Research Fellow (CR) and R.H. is a senior Research Associate (MR) of the F.R.S. – FNRS.

## Authors contribution

A.P. and R.H. conceived and designed the experiments. A. B-V performed the virulence assays in *Galleria mellonella* (Fig. 5 and Fig. S6). A.P. performed all the other experiments presented in this manuscript. A.P. and R.H. wrote the manuscript.

## Supplemented Figure legends

**Figure S1.**
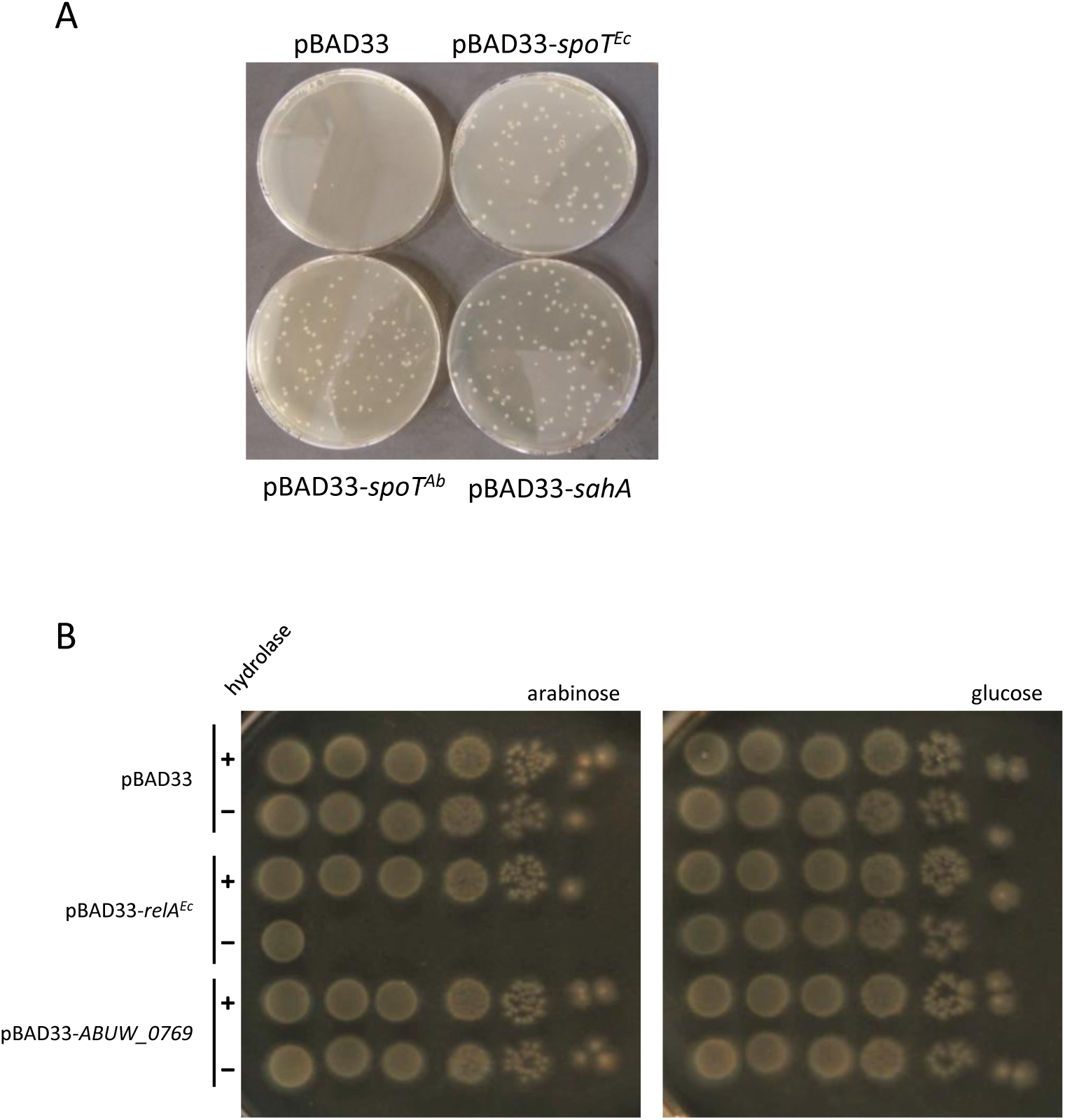
SahA can hydrolyze (p)ppGpp *in vivo* in *E. coli* while putative SAS does not seem to produce (p)ppGpp in *E. coli*. (**A**) Growth on LB plates supplemented with Kanamycin and Arabinose after P1 mediated transduction of a *spoT*::*kan^R^* lysate into *E. coli* MG1655 expressing *spoT^EC^*, *spoT^AB^* or *sahA* from a pBAD33 vector. (**B**) Viability of *E. coli* MG1655 WT (hydrolase +) or Δ*relA* Δ*spoT* (hydrolase -) strains expressing *relA^EC^* or *ABUW_0769* (putative SAS) from a pBAD33 vector. Overnight cultures were serial diluted (1:10), spotted on LB plates with 0.5% of glucose or arabinose and incubated overnight at 37 °C

**Figure S2.**
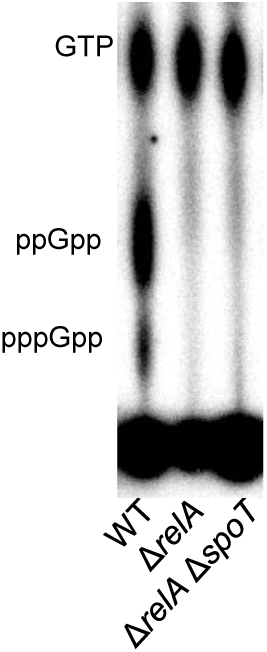
RelA seems to be the sole (p)ppGpp synthetase in *A. baumannii*. Nucleotides extracted from cells treated with serine hydroxamate (SHX) for 15min were analyzed on thin layer chromatography. *A. baumannii* AB5075 WT, Δ*relA*, Δ*relA* Δ*spoT* are represented.

**Figure S3.**
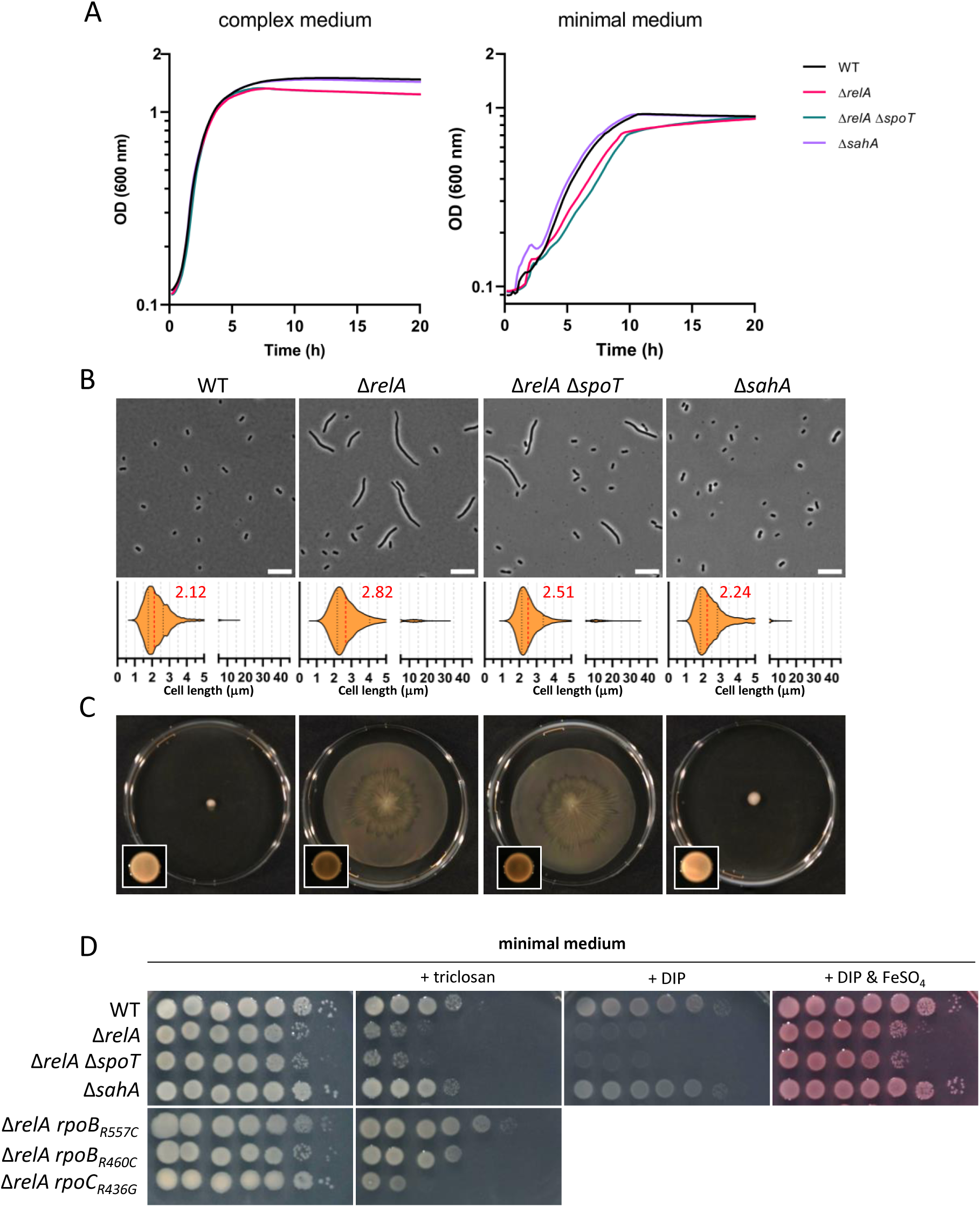
The Δ*sahA* and Δ*relA* Δ*spoT* mutants show similar phenotypes compared to the WT and Δ*relA* strains, respectively. **(A)** Representative growth of WT, Δ*relA*, Δ*relA* Δ*spoT* and Δ*sahA* at 37 °C in complex (LB) or minimal synthetic (MOPSX) medium for 20 h in 96-well plates. **(B)** Microscopic analysis of WT, Δ*relA*, Δ*relA* Δ*spoT* and Δ*sahA* cells grown overnight at 37 °C in liquid LB complex medium. A representative picture (scale bar = 10 µm) is shown for each strain accompanied by a violin plot showing cell size distribution; the median size (µm) is indicated in red. **(C)** Surface motility of cells described in (B) on low agar (0.5%) LB complex medium. The appearance and the color of the colonies can also be observed on the picture in the box in bottom left corners. Pictures for both experiments were taken after 24h incubation at 37 °C. **(D)** Viability of WT, Δ*relA*, Δ*relA* Δ*spoT,* Δ*sahA, rpoB_R557C_*, Δ*relA rpoB_R460C_* and Δ*relA rpoB_R436G_* strains on minimal synthetic medium (M9X) with or without supplementation [triclosan (0.125 µM); DIP (150 µM); FeSO_4_ (50 µM)]. Overnight cultures were serial diluted (1:10), spotted on plates and incubated overnight at 37 °C.

**Figure S4.**
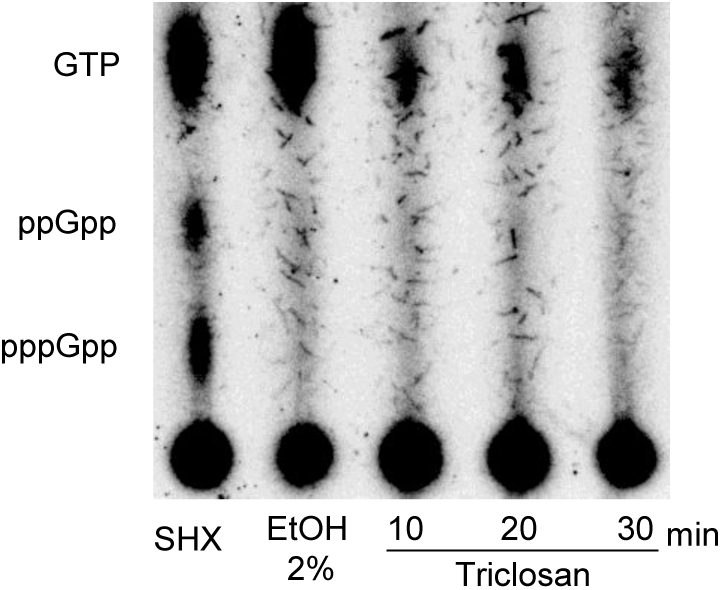
Triclosan does not induce (p)ppGpp production *A. baumannii* AB5075. Nucleotides extracted from WT cells exposed to either serine hydroxamate (SHX) or ethanol (EtOH) for 10 and 30 min respectively or Triclosan (250 µg/mL) were analyzed on thin layer chromatography.

**Figure S5.**
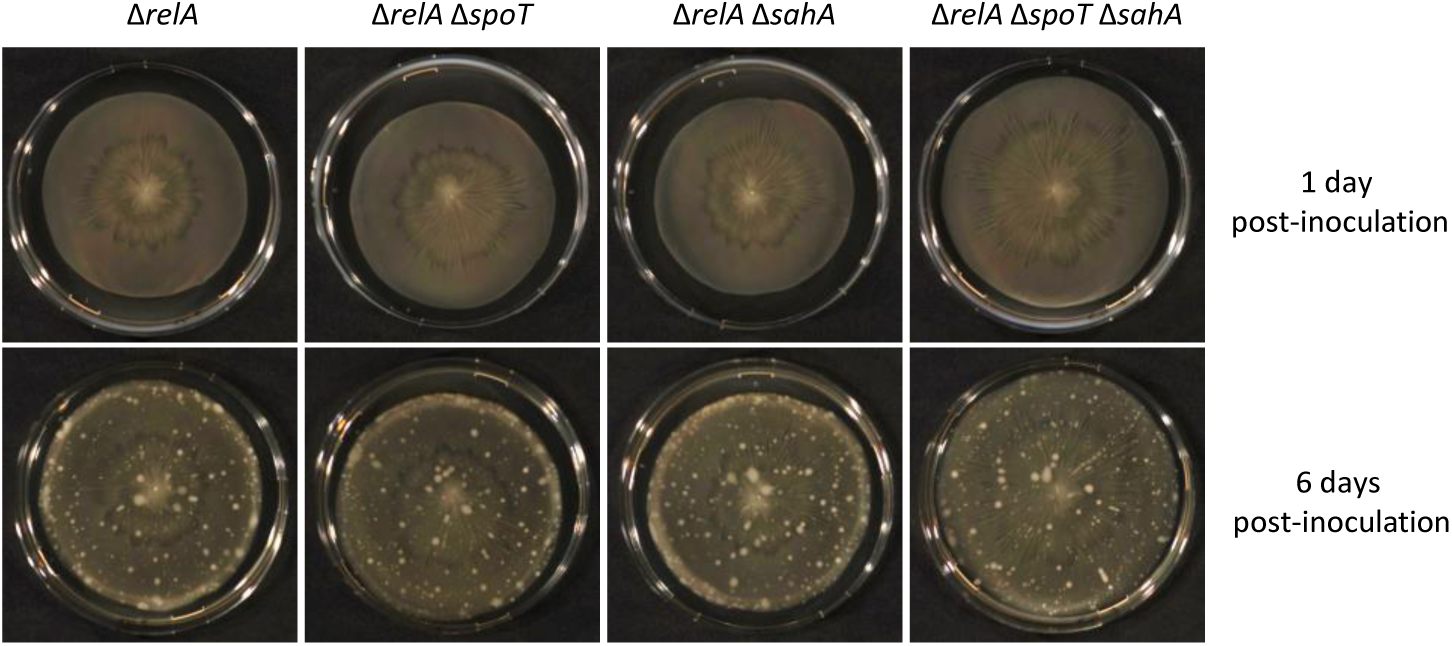
Putative motility suppressors easily selected on low agar medium in (p)ppGpp^0^ backgrounds. The WT, Δ*relA*, Δ*relA* Δ*spoT* and Δ*relA* Δ*spoT* Δ*sahA* strains were inoculated on low agar medium. After 6 days at 37 °C, white non-motile colonies growing on top of the motility lawn can be observed.

**Figure S6.**
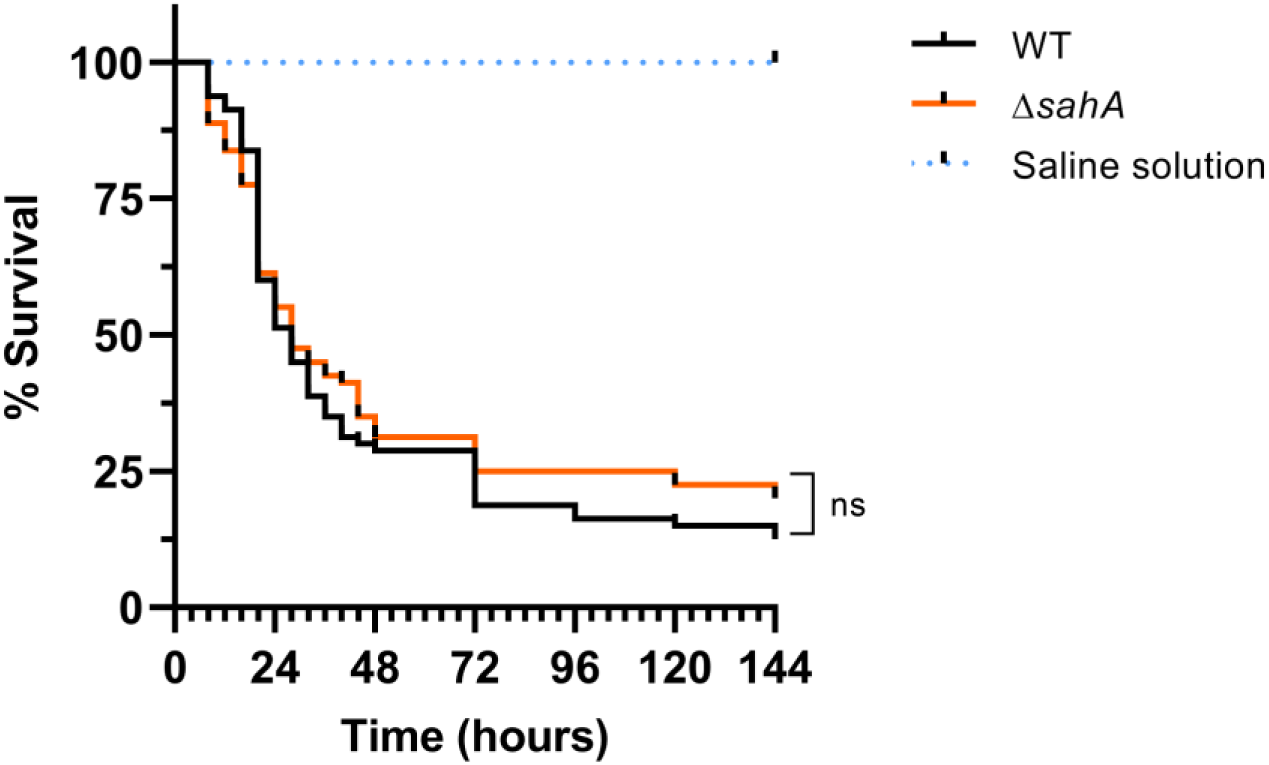
The small alarmone hydrolase *sahA* is not required for the virulence in the *Galleria mellonella* model. Kaplan-Meier survival analysis of *Galleria mellonella* larvae infected with the WT and Δ*sahA* strains. Each curve represents the average of three independent experiment for a total of 80 larvae per condition. No significant differences between the survival curve of Δ*sahA* mutant and the WT one (Log-rank (Mantel-Cox) test; *p* = 0.4022).

**Figure S7.**
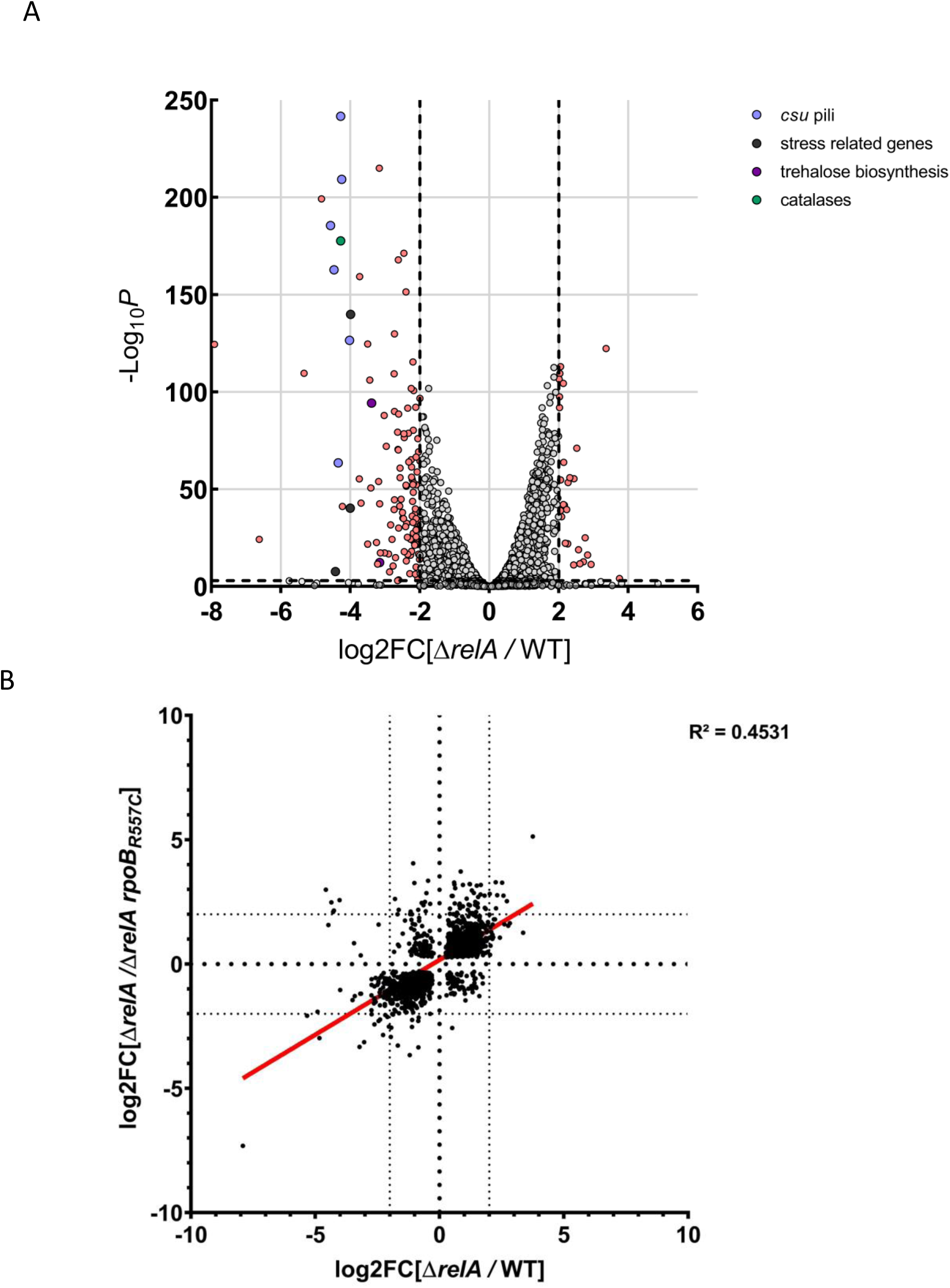
Lack of (p)ppGpp during exponential growth phase in complete medium has only on mild effect on the transcriptome profile. **(A)** Volcano plot representing the genes differentially expressed in the Δ*relA* mutant compared to the WT strain on liquid complete medium. Genes with a |log_2_FC| ≥ 2 and –Log_10_ *p*-value ≥ 3 are represented in light pink or with a color code for group of genes associated with known functions (blue: csu pili; black: stress related genes; purple: trehalose biosynthesis; dark green: catalases). **(B)** xy-plot representing the differential expression of genes between the Δ*relA*/WT *vs* Δ*relA/*Δ*relA rpoB_R557C_* in liquid complex medium. Each dot represents the log_2_FC value for the strains indicated under brackets (only genes with FDR-adjusted *p* ≤ 0.005 were kept).

